# Alcohol-Induced Increase in the Content of CYP2E1 in Human Liver Microsomes Causes a Multifold Activation of CYP3A4 and Attenuates its Cooperativity

**DOI:** 10.1101/2020.03.09.984211

**Authors:** Bikash Dangi, Nadezhda Y. Davydova, Nikita E. Vavilov, Victor G. Zgoda, Dmitri R. Davydov

**Author notes:** **Corresponding author:** Dmitri R. Davydov, Department of Chemistry, Washington State University, MS 4630, Pullman, WA 99164-4630.

## Abstract

Here we investigate the effect of alcohol-induced increase in the content of CYP2E1 in human liver microsomes (HLM) on the function of CYP3A4. In these studies we used a model that implements enrichment of HLM samples with CYP2E1 through membrane incorporation of the purified protein. Enrichment of HLM with CYP2E1 considerably increases the rate of metabolism of 7-benzyloxyquinoline (BQ) and attenuates the homotropic cooperativity observed with this CYP3A4-specific substrate. Incorporation of CYP2E1 also eliminates the activating effect of α-Naphthoflavone (ANF) on BQ metabolism seen in some untreated HLM samples. To probe the physiological relevance of these effects we compared three pooled preparations of HLM from normal donors (HLM-N) with a preparation obtained from heavy alcohol consumers (HLM-A). The composition of the P450 pool in all four samples was characterized with mass-spectrometric determination of 11 cytochrome P450 species. The molar content of CYP2E1 in HLM-A was from 2.5 to 3.3 times higher than that found in HLM-N. In contrast, the content of CYP3A4 in HLM-A was the lowest among all four HLM samples. Despite of that, HLM-A exhibited much higher rate of metabolism and lower degree of homotropic cooperativity with BQ, similar to that observed in CYP2E1-enriched HLM-N. In order to substantiate the hypothesis on the involvement of physical interactions between CYP2E1 and CYP3A4 in the observed effects we probed hetero-association of these proteins in Supersomes™ containing recombinant CYP3A4 with a technique based on homo-FRET and employing CYP2E1 labeled with BODIPY-618 maleimide. These experiments demonstrated high affinity interactions between the two enzymes and revealed an inhibitory effect of ANF on their hetero-association. Our results demonstrate that the catalytic activity and allosteric properties of CYP3A4 are fundamentally dependent on the composition of the cytochrome P450 ensemble and imply a profound impact of chronic alcohol exposure on the pharmacokinetics of drugs metabolized by CYP3A4.

## Introduction

The core of the drug-metabolizing ensemble in human liver is the ensemble of multiple P450 species co-localized in the membrane of endoplasmic reticulum (ER). There is an emerging recognition of the fact that multiple P450 enzymes interact with each other with the formation of heteromeric complexes where the functional properties of individual P450 enzymes are largely modified (Davydov, 2011; Reed and Backes, 2012; Davydov, 2016; Reed and Backes, 2016). Resent findings point out the association of dissimilar P450s as the major cause for poor correlation between the composition of the P450 ensemble and its drug metabolizing profile demonstrated with studies in a large set of normal human liver microsomes (Gao et al., 2016; Zhang et al., 2016).

Of particular importance are the changes in P450-P450 crosstalk induced by alcohol consumption. Although the multi-fold increase in the content of CYP2E1 in liver observed in alcohol consumers is well documented (Cederbaum, 1998; Dupont et al., 1998; Lieber, 1999; Cederbaum, 2006), the involvement of CYP2E1 in alcohol-drug interactions is commonly considered insignificant due to a minor role of this enzyme in drug metabolism (Jang and Harris, 2007). However, the effects of induction of CYP2E1 may stretch beyond the changes in CYP2E1-dependent drug metabolism and involve the effects of CYP2E1 on the functional properties of other drug-metabolizing enzymes.

In order to probe the crosstalk between CYP2E1 and other P450 species and elucidate its role in alcohol-drug interactions we established a model of alcohol-induced increase in CYP2E1 content that implements enrichment of HLM samples with CYP2E1 through membrane incorporation of the purified protein (Davydov et al., 2015; Davydov et al., 2017; Davydova et al., 2019). We demonstrated that the adopted CYP2E1 becomes a fully-functional member of the drug-metabolizing ensemble and interacts with other P450 enzymes with the formation of heteromeric complexes (Davydova et al., 2019). Studying the effect of enrichment of HLM with CYP2E1 on the function of other P450 enzymes we demonstrated a CYP2E1-induced activation of CYP1A2 and the associated re-routing of the metabolism of 7-ethoxy-4-cyanocoumarin (CEC), the substrate concurrently metabolized by CYP2C19 and CYP1A2, towards the latter enzyme (Davydova et al., 2019).

The present study continues our efforts on revealing the impact of alcohol-induced increase in CYP2E1 content on drug metabolism. Here we explore the effects of CYP2E1 on the functional properties of CYP3A4, the enzyme that metabolizes about 50% of drugs on the market (Guengerich, 1999). As a probe substrate we used 7-benzyloxyquinoline (BQ), a fluorogenic substrate highly selective for CYP3A4 (Stresser et al., 2002). In addition to studying the effect of CYP2E1 on the parameters of BQ metabolism in a series of microsomal preparations with thoroughly characterized composition of the P450 pool, we also probed its impact on the effect of *α*-naphthoflavone, a prototypical allosteric effector of CYP3A4. In order to assess the relevance of the results obtained with CYP2E1-enriched microsomes to the changes caused by the effects of chronic alcohol exposure we also compared the parameters of BQ metabolism in a pooled HLM preparation obtained from 10 alcoholic donors with those exhibited by three pooled HLM preparations obtained from donors with no history of alcohol dependence. Our results demonstrate that the catalytic activity and allosteric properties of CYP3A4 are fundamentally dependent on the composition of the cytochrome P450 ensemble in human liver and imply a profound impact of chronic alcohol exposure on the pharmacokinetics of drugs metabolized by CYP3A4.

Furthermore, probing possible mechanisms of CYP2E1-dependent activation of CYP3A4 revealed in this study we investigated the interactions between the two proteins in the microsomal membrane using the technique based on homo-FRET in the oligomers of CYP2E1 labeled with BODYPY-618 maleimide (CYP2E1-BODIPY) (Davydova et al., 2019). High-affinity interactions between CYP2E1-BODIPY with CYP3A4 and their modulation by *α*-naphthoflavone revealed in these studies suggest a direct involvement of hetero-association between CYP2E1 and CYP3A4 in the observed effects.

## Materials and methods

### Chemicals

7-benzyloxyquinoline (BQ) was a product of CypEx (Dundee, UK). 7-hydroxycoumarin was obtained from Acros Organics, a part of Thermo Fisher Scientific (New Jersey, NJ). 7,8-benzoflavone (*α*-naphthoflavone, ANF) was a product of Indofine Chemical Company (Hillsborough, NJ). All other reagents were of ACS grade and used without additional purification.

### Protein expression and purification

N-terminally truncated (*Δ*3-20) and C-terminally His-tagged CYP2E1 (Spatzenegger et al., 2003) was expressed in *E. coli* TOPP3 cells and purified as described earlier (Davydova et al., 2019).

### Pooled human liver microsomes and their characterization with mass-spectrometry

The preparation of Human Liver Microsomes (HLM) obtained from 10 donors (mixed gender) with a history of chronic alcohol exposure (lot FVT) was purchased from BioIVT corporation (Baltimore, MD). This preparation is referred hereinafter as HLM-A. We also studied three different preparations of pooled human liver microsomes from 50 donors (mixed gender) without reported history of alcohol exposure. The preparation obtained by differential centrifugation of pooled human liver S9 fraction (the product of BD Gentest, lot number 3212595) is referred to as HLM-N1. The HLM samples referred here as HLM-N2 and HLM-N3 are InVitroCYP™ M-class 50-donor mixed gender pooled HLM preparations, lots LBA and LFJ respectively obtained from BioIVT corporation (Baltimore, MD).

The composition of the cytochrom P450 ensemble in these preparations was characterized by mass-spectrometric analysis with a triple quadrupole mass spectrometer using the method of multiple reaction monitoring (MRM) as described previously (Davydova et al., 2019). Content of NADPH-cytochrome P450 reductase and cytochromes P450 1A2, 2A6, 2B6, 2D6, 2C8, 2C9, 2C18, 2C19, 2E1, 3A4 and 3A5 was determined based on the concentration of protein-specific tryptic peptides in trypsin-digested samples with the use of the stable isotope labeled internal standards (SIS) (Davydova et al., 2019). For each protein, one standard peptide with three transitions was used. The peptides, which sequences are provided in Table 1, were arranged into one Selected Reaction Monitoring (SRM) assay. Mass-spectrometry measurements of P450 content were performed using the equipment of “Human Proteome” Core Facilities of the Institute of Biomedical Chemistry (Moscow, Russia).

**Table 1.**
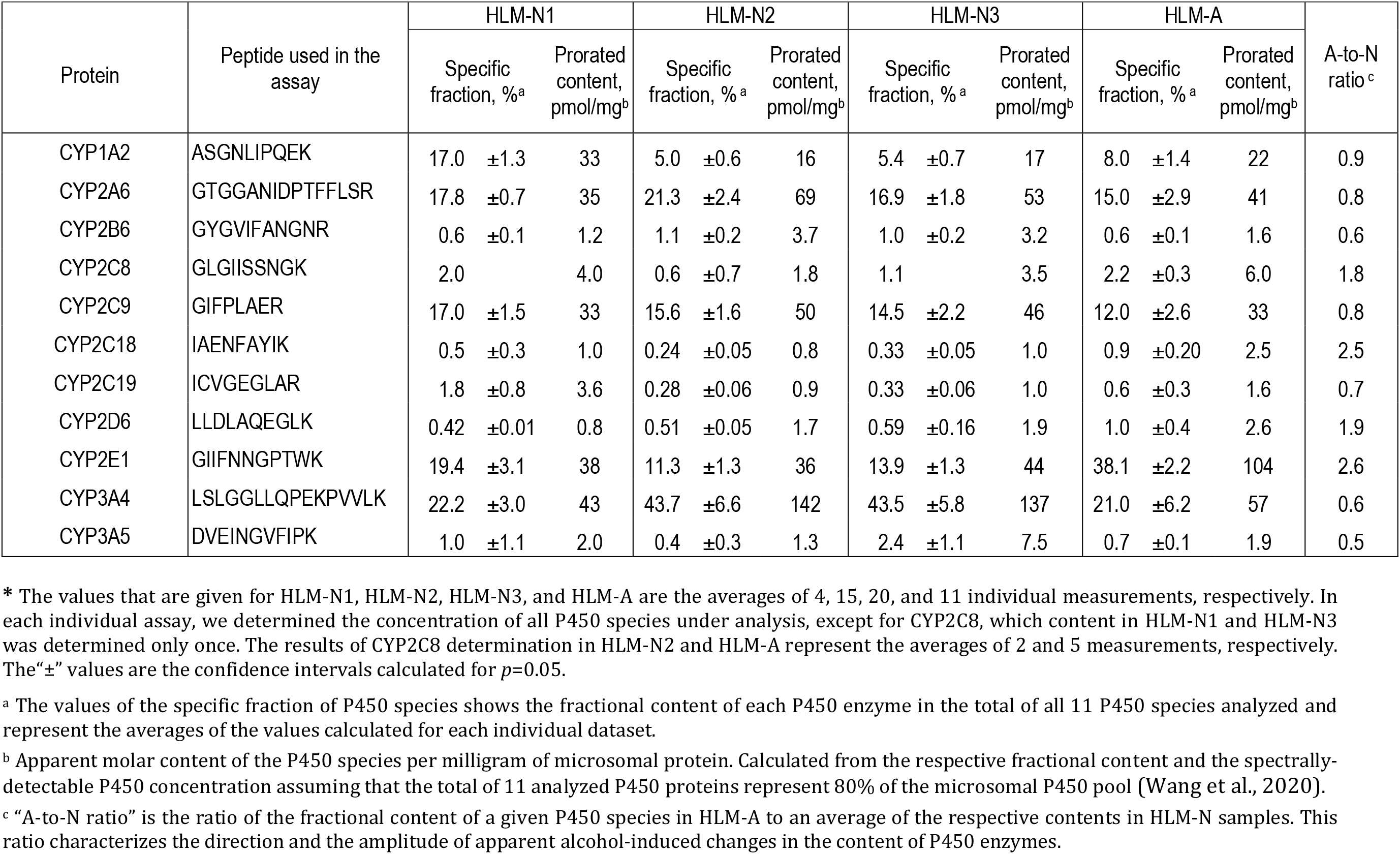
Composition of the cytochrome P450 pool in HLM preparations under study*.

### Characterization of the content of protein, phospholipids, NADPH-cytochrome P450 reductase and cytochromes P450 in HLM

Determinations of protein and phospholipid concentrations in microsomal suspensions were performed with the bicinchoninic acid procedure (Smith et al., 1985) and through the determination of total phosphorus in a chloroform/methanol extract according to Bartlett (Bartlett, 1959), respectively. The concentration of NADPH-cytochrome P450 reductase in microsomal membranes was determined based on the rate of NADPH-dependent reduction of cytochrome *c* at 25 °C and the effective molar concentration of CPR was estimated using the turnover number of 3750 min^-1^ (Davydova et al., 2019). The total concentration of cytochromes P450 in HLM was determined with a variant of “oxidized CO versus reduced CO difference spectrum” method as described earlier (Davydova et al., 2019).

### Preparation of CYP2E1-enriched HLM samples

Incorporation of CYP2E1 into HLM was performed by incubation of undiluted suspensions of HLM (20-25 mg/ml protein, 10-13 mM phospholipid) in 125 mM K-Phosphate buffer containing 0.25M Sucrose with purified CYP2E1 for 16 - 20 hours at 4°C at continuous stirring. CYP2E1 was added in the amount ranging from 0.25 to 1 molar equivalent to the endogenous cytochrome P450 present in HLM. Following the incubation, the suspension was diluted 4-8 times with 125 mM K-Phosphate buffer, pH 7.4 containing 0.25 M sucrose and centrifuged at 53,000 rpm (150,000 g) in an Optima TLX ultracentrifuge (Beckman Coulter Inc., Brea, CA, USA) with a TLA100.3 rotor for 90 min at 4 °C. The pellet was resuspended in the same buffer to the protein concentration of 15-20 mg/ml. The amount of incorporated cytochrome P450 was calculated from the difference between the heme protein added to the incubation media and the enzyme found in the supernatant. According to the results of this assay, our procedure resulted in incorporation of 96 - 98% of the added protein into the microsomal membrane.

### Fluorimetric assays of BQ metabolism

The rate of debenzylation of 7-benzyloxyquinoline was measured with a real-time continuous fluorometric assay using a Cary Eclipse fluorometer (Agilent Technologies, Santa Clara, CA, USA) or a custom-modified PTI QM-1 fluorometer (Photon Technology International, New Brunswick, NJ (Davydov et al., 2017). In the experiments with Cary Eclipse the excitation was performed with a monochromatic light centered at 405 nm with 5 nm bandwidth. In the case of PTI QM-1 the excitation light centered at 405 nm was emitted by a CPS405 collimated laser diode module (Thorlabs Inc, Newton, NJ). The emission wavelength was set at 516 nm with a 20 nm slit. The rate of formation of 7-hydroxyquinolyne was estimated by determining the slope of the linear part of the kinetic curve recorded over a period of 2 - 3 min.

All kinetic assays were performed in 0.1 M Na-HEPES buffer, pH 7.4, containing 60 mM KCl. In the case of the use of Cary Eclipse the total volume of the incubation mixture was equal to 300μl and a 5 × 5 mm quartz cell was used. In the experiments with PTI QM-1 fluorometer we used a 3 × 3 mm quartz cell and the volume of the sample was equal to 60μl. With both instruments the kinetic assays were carried out at continuous stirring and the temperature was maintained at 30 °C with a circulating water bath. An aliquot of 15-24 mM stock solution of BQ in acetone was added to attain the desired substrate concentration in the range of 0.5 - 250 μM. The reaction was initiated by addition of 20 mM solution of NADPH to the concentration of 200 μM. Fitting of the dependencies of the reaction rate on the substrate concentration to Michaelis-Menten and Hill equations was performed with a combination of Marquardt and Nelder-Mead non-linear regression algorithms as implemented in our SpectraLab software (Davydov et al., 1995; Davydov et al., 2016).

### Monitoring the interactions of CYP2E1-BODIPY with microsomal membranes

Labeling of CYP2E1 with BODIPY-618 maleimide was performed at 2:1 label to protein ratio as described earlier (Davydov et al., 2015; Davydov et al., 2017; Davydova et al., 2019). The studies of interactions of CYP2E1-BODIPY with microsomal membranes were performed with the use of a Cary Eclipse spectrofluorometer equipped with a Peltier 4-position cell holder. The excitation of donor phosphorescence was performed with monochromatic light centered at 405 with 20 nm bandwidth. Alternatively the measurements were done with the use a custom-modified PTI QM-1 fluorometer (Photon Technology International, New Brunswick, NJ) equipped with a thermostated cell holder and a refrigerated circulating bath. In this case the excitation was performed at 405 nm with a CPS405 collimated laser diode module (Thorlabs Inc, Newton, NJ). The spectra in the 570 - 750 nm wavelength region were recorded repetitively with the time interval varying from 0.5 to 15 min during the course of monitoring (5 - 16 hours). The experiments were performed at continuous stirring at 4 °C in 100 mM Na-Hepes buffer (pH 7.4) containing 150 mM KCl and 250 mM sucrose.

Calculations of the surface density of cytochromes P450 in microsomal membranes (C_CYP2E1_) were based on the molar ratio of membranous phospholipids to incorporated CYP2E1 (*R*_L/P_). For our calculations we assumed the area of microsomal membrane corresponding to one molecule of phospholipid in a monolayer to be equal to 0.95 nm^2^ (Wibo et al., 1971), similar to the approach used in our earlier reports (Davydov et al., 2015; Davydov et al., 2017; Davydova et al., 2019). The use of this value results in the following relationship between C_CYP2E1_ and the *R*_L/P_:

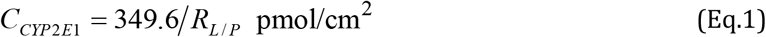

Analysis of series of spectra obtained in fluorescence spectroscopy experiments was done by principal component analysis (PCA) (Davydov et al., 1995) as previously described (Davydova et al., 2019). The equation for the equilibrium of binary association (dimerization) used in the fitting of oligomerization isotherms (dependencies of FRET efficiency on the concentration of P450 in membranes) had the following form:

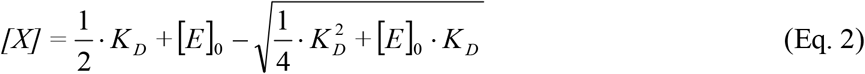

where [*E*]_0_, [*X*], and *K*_D_ are the total concentration of the associating compound (enzyme), the concentration of its dimers, and the apparent dissociation constant, respectively. In order to use this equation in fitting of the dependencies of the relative increase in fluorescence observed at enzyme concentration [*E*]_0_ (*R*_E_) equation (1) was complemented with the parameter *R*_max_. This parameter corresponds to the value of *R*_E_ observed upon a thorough dissociation of completely oligomerized enzyme:

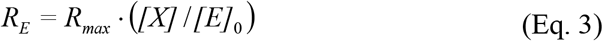

The parameter *R*_max_ is determined by the efficiency of FRET (*E*_FRET_) according to the following relationship:

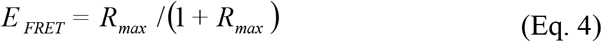

Fitting of the titration isotherms to the above equations was performed with non-linear regression using a combination of Nelder-Mead and Marquardt algorithms as implemented in our SpectraLab software (Davydov et al., 1995).

## Results

### Characterization of the composition of the cytochrome P450 pool in HLM samples

The preparations of HLM used in this study were characterized with mass-spectrometry by determining the content of NADPH-cytochrome P450 reductase, cytochrome *b*_5_ and 11 major cytochrome P450 species, namely CYP1A2, CYP2A6, CYP2B6, CYP2C8, CYP2C9, CYP2C18, CYP2C19, CYP2E1, CYP3A4, and CYP3A5. The results of our analysis of the composition of the P450 pool in HLM are summarized in Table 1.

The total content of cytochromes P450 and the concentration of NADPH-cytochrome reductase (CPR) and cytochrome *b*_5_ were also quantified with absorbance spectroscopy and determination of the rate of NADPH-dependent reduction of cytochrome *c*. In Table 2 we compare the results of spectrophotometric and LC-MS/MS assays of the contents of CPR, cytochrome *b*_5_ and cytochrome P450. Significant scatter in the degrees of coverage of the concentrations determined in the spectrophotometric assays by the values derived from the LC-MS/MS analysis suggest a substantial variability in the efficiency of proteolytic digestion in our experiments. Due to this variability, the absolute quantities of the cytochrome P450 species presented in Table 1 were calculated from the respective fractional contents and the amount of spectrally-detectable P450 hemeprotein. In these calculations we assumed that the total of 11 analyzed P450 proteins represent 80% of the microsomal P450 pool. This estimate was calculated from the data Wang and co-authors, who determined the content of 25 P450 species in a series of 102 individual HLM samples (Wang et al., 2020). The values shown in Table 1 for HLM-N1, HLM-N2, HLM-N3, and HLM-A are the mean values averaged over the results of 4, 15, 20, and 11 individual measurements, respectively. As expected, the fractional content of CYP2E1 in HLM-A was from 2.7 to 3.5 fold higher than that detected in the HLM-N samples.

**Table 2.** Contents of NADPH-cytochrome P450 reductase, cytochrome *b*_5_ and cytochromes P450 and in the microsomal preparations under study.

| Protein | Parameter | HLM-N1 | HLM-N2 | HLM-N3 | HLM-A |
| --- | --- | --- | --- | --- | --- |
| NADPH-cytochrome P450 reductase (CPR) <sup>a</sup> | Content based on activity assays, pmol/mg | 58 ±10 | 63 ±12 | 51 ±12 | 35 ±4 |
|  | Content based on LC-MS/MS assays, pmol/mg | 30 ±23 | 43 ±11 | 42 ±9 | 44 ±8 |
|  | LC-MS/MS coverage, % <sup>c</sup> | 52 ±40 | 68 ±17 | 82 ±20 | 127 ±23 |
| Cytochrome *b*<sub>5</sub> <sup>a</sup> | Content based on UV-VIS absorbance assays, pmol/mg | 375 ±19 | 288 ±5 | 329 ±2 | 250 ±30 |
|  | Content based on LC-MS/MS assays, pmol/mg | 117 ±64 | 163 ±72 | 164 ±48 | 23 ±8 |
|  | LC-MS/MS coverage, % <sup>c</sup> | 31 ±17 | 57 ±25 | 44 ±17 | 9 ±1 |
| Cytochrome P450 <sup>b</sup> | Content based on UV-VIS absorbance assays, pmol/mg | 244 ±20 | 405 ±95 | 394 ±53 | 341 ±70 |
|  | Content based on LC-MS/MS assays, pmol/mg | 113 ±67 | 355 ±135 | 323 ±94 | 61 ±25 |
|  | LC-MS/MS coverage, % <sup>c</sup> | 46 ±27 | 88 ±33 | 82 ±24 | 18 ±7 |

\* The values given for the assays based on absorbance (UV-VIS) spectroscopy are the averages of 3 – 6 individual measurements. The results of LC-MS/MS assays with HLM-N1, HLM-N2, HLM-N3, and HLM-A are represented by the averages of 4, 15, 20, and 11 individual measurements, respectively. The “±” values are the confidence intervals calculated for *p*=0.05.
<sup>a</sup> The contents of CPR and cytochrome *b*<sub>5</sub> determined with LC-MS/MS are based on the detected amount of tryptic peptides NPFLAAVTTNR and STWLILHHK, respectively.
<sup>b</sup> The values of the apparent molar content of cytochromes P450 determined by LC-MS/MS represent the totals of the recovered amounts of 11 analyzed cytochrome P450 species.
<sup>c</sup> The coverage represent the percent fraction of the amount of proteins recovered with LC-MS/MS from their amounts determined with the absorbance spectroscopy assays.

### Parameters of BQ metabolism exhibited by four HLM-preparations

In order to probe the effect of the composition of the P450 ensemble on the functional properties of CYP3A4 we determined the parameters of 0-debenzylation of BQ by all four microsomal preparations under study. We also probed the effect of ANF, the prototypical allosteric effector of the enzyme. Results of these studies are summarized in Table 3 and illustrated in Fig. 1. It should be noted that the rate of metabolism of BQ in HLM in this study was normalized on the concentration of CPR in the microsomal membrane (see Materials and Methods). In view of a high excess of cytochromes P450 over CPR in HLM, this normalization was considered as the most appropriate approach for comparing the rates of metabolism in the HLM preparations with different composition of the cytochrome P450 ensemble.

**Table 3.** Parameters of BQ metabolism obtained with four HLM preparations and their modulation by *α*-naphthoflavone*.

| HLM preparation<sup>a</sup> | S<sub>50</sub>, μM | | N | | V<sub>max</sub>, min<sup>-1</sup> | |
| --- | --- | --- | --- | --- | --- | --- |
|  | No effector | +ANF | No effector | +ANF | No effector | +ANF |
| HLM-N1 | 40.2 ±12.2 | **10.7 ±4.8 (0.012)** | 2.24 ±0.37 | **0.83 ±0.12 (0.002)** | 16.4 ±1.8 | **24.9 ±2.8 (0.002)** |
| HLM-N1+2E1 | **89.2 ±20.4 (0.012)** | **12.4 ±4.3 (0.012)** | **1.26 ±0.12 (0.009)** | **0.87 ±0.16 (0.020)** | **31.9 ±5.0 (<10<sup>-3</sup>)** | **19.9 ±2.7 (0.014)** |
| HLM-N2 | 36.7 ±10.3 | **7.4 ±3.8 (0.006)** | 2.09 ±0.44 | **1.22 ±0.30 (0.033)** | 71.4 ±11.4 | **32.6 ±3.5 (0.003)** |
| HLM-N2+2E1 | **89.1 ±34.3 (0.031)** | **12.0 ±4.7 (0.014)** | **1.28 ±0.09 (0.011)** | 1.02 ±0.24 (0.068) | 83.7 ±38.8 (0.573) | 46.1 ±11.1 (0.131) |
| HLM-N3 | 63.3 ±3.2 | **19.7 ±6.7 (<10<sup>-3</sup>)** | 1.92 ±0.02 | **1.02 ±0.37 (0.010)** | 45.7 ±5.1 | 49.3 ±7.9 (0.528) |
| HLM-N3+2E1 | 107 ±55 (0.248) | **16.3 ±6.9 (0.019)** | **1.13 ±0.03 (<10<sup>-6</sup>)** | **0.86 ±0.08 (0.003)** | **77.3 ±4.9 (0.001)** | 46.8 ±16.6 (0.082) |
| HLM-A | 59.6 ±12.5 | **25.0 ±10.7 (0.006)** | 1.02 ±0.15 | **0.99 ±0.20 (0.025)** | 101 ±20 | 113 ±24 (0.934) |
| HLM-A+2E1 | 65.4 ±25.5 (0.665) | **23.0 ±13.1 (0.044)** | 1.47 ±0.30 (0.295) | **0.85 ±0.26 (0.033)** | **147 ±30 (0.028)** | 127 ±4 (0.318) |

\* The values given in the table are the averages of 3 - 7 individual measurements. The“ ± ” values are the confidence intervals calculated for $p=0.05$ . The values in parentheses represent the $p$ -values of Student’s t-test for the hypothesis of equality of the respective values with the value obtained with untreated microsomes (for the values in “No effector” columns) or at no ANF added (for the values in “+ANF” columns). The values characterized with $p$ -values $\leq 0.05$ are shown in bold to emphasize the effects with high statistical significance.
<sup>a</sup> The CYP2E1-enriched samples represented in this table are the samples where the content of added CYP2E1 was equal to the total P450 content in untreated HLM.

**Fig. 1.**
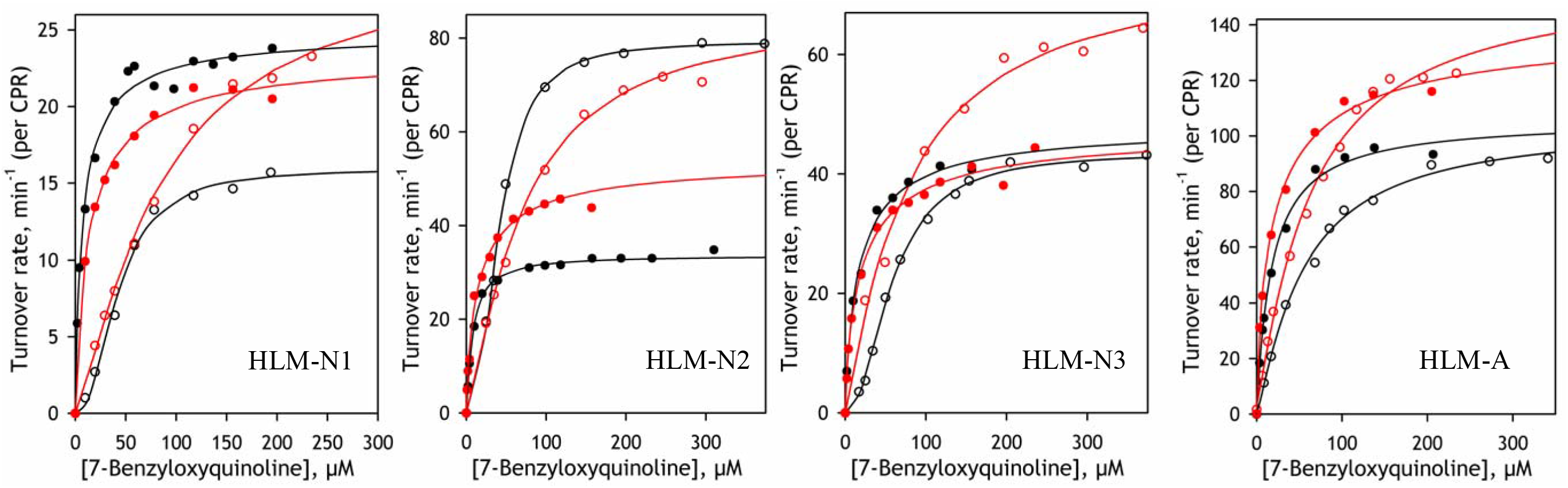
BQ metabolism in intact and CYP2E1-enriched HLM samples. The figure show the dependencies of the rate of BQ turnover on substrate concentration in intact (black) and CYP2E1-enriched (red) HLM preparations. The dependencies obtained in the absence of added effector are shown in open circles, while the data obtained in the presence of 25 μM ANF are designated by closed circles. The CYP2E1-enriched samples exemplified in this figure are the samples where the content of added CYP2E1 was equal to the total P450 content in HLM (“1:1 incorporation”). The datasets shown on this figure were obtained by averaging the results of 3 - 5 individual experiments. The solid lines show the results of fitting of the datasets with the Hill equatrion.

In all three samples of HLM-N the metabolism of BQ revealed a prominent homotropic cooperativity characterized with the Hill coefficient of 1.9 - 2.2 (Fig. 1, curves shown in black open circles), in a good agreement with the previous reports (Stresser et al., 2002; Davydov et al., 2008). In HLM-N samples the maximal rate of reaction increases concomitant with increasing fractional content of CYP3A4 in microsomes (Table 3). Strikingly, despite the low content of CYP3A4 in HLM-A, this preparation is characterized with the highest maximal rate of BQ metabolism over all four HLM samples studied. *V*_max_ value exhibited by HLM-A is over 6 times higher than the value obtained with HLM-N1, where the fractional content of CYP3A4 (28%) is comparable to that observed in HLM-A (21%). The values of *V*_max_ obtained with HLM-N2 and HLM-N3, where the fractional content of CYP3A4 is around 40%, are respectively 1.4 and 2.2 times lower than that characteristic to HLM-A. This multifold activation of BQ metabolism in HLM-A is associated with the attenuation of homotropic cooperativity, which is revealed in a considerable decrease of the Hill coefficient (Table 3).

The contrasting difference between HLM-A and HLM-N preparation in the rate of BQ metabolism is illustrated in Fig. 3, where the *V*_max_ values are plotted against the molar ratio of CYP3A4 to CPR determined in these preparations with LC-MS/MS. As it might be expected, the *V*_max_ value observed with HLM-N preparations and calculated per molar content of CPR displays an evident proportionality to the content of CYP3A4A (Fig. 2a). However, the rate observed with HLM-A, where the content of CYP3A4 is the lowest out of all four preparations, deviates from this proportionality being the highest of all four values under comparison. The difference between HLM-A and HLM-N becomes even more contrasting when the rates of turnover are calculated per molar content of CYP3A4. 0bviously, this way of normalization attenuates the difference between the three HLM-N samples, whereas the turnover number calculated for HLM-A becomes almost an order of magnitude higher than the values calculated for HLM-N samples (Fig. 2b).

**Fig. 2.**
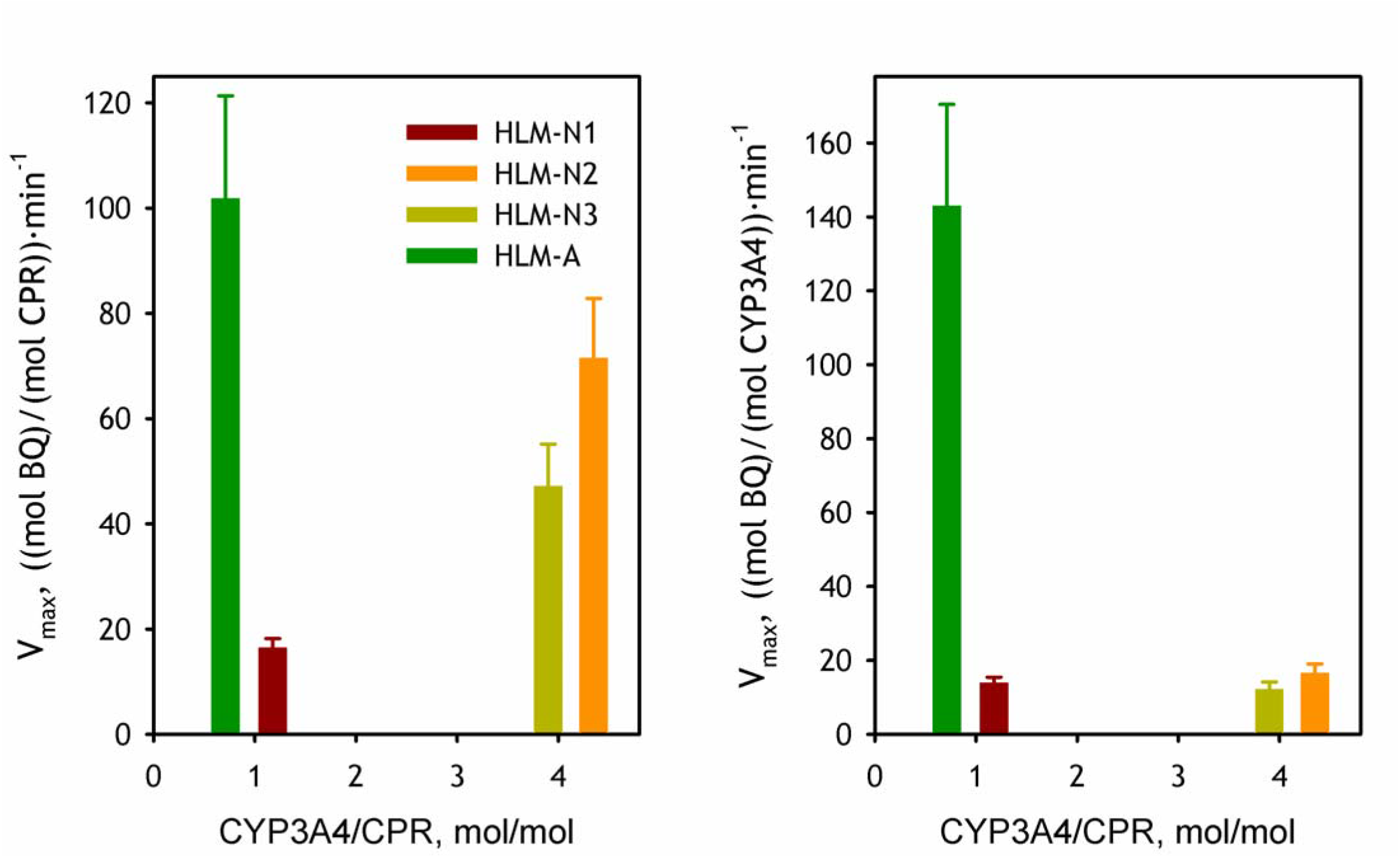
The relationship between the CYP3A4 content in HLM preparations and the rate of BQ metabolism an no added effector. The maximal velocity of BQ debenzylation is plotted against the molar ratio of CYP3A4 to CPR in microsomal membranes calculated from the results of LC-MS/MS analysis (Table 1). Left panel shows the *V*_max_ values normalized on the concentration of CPR, while the right panel shows the same results normalized per CYP3A4 content.

**Fig. 3.**
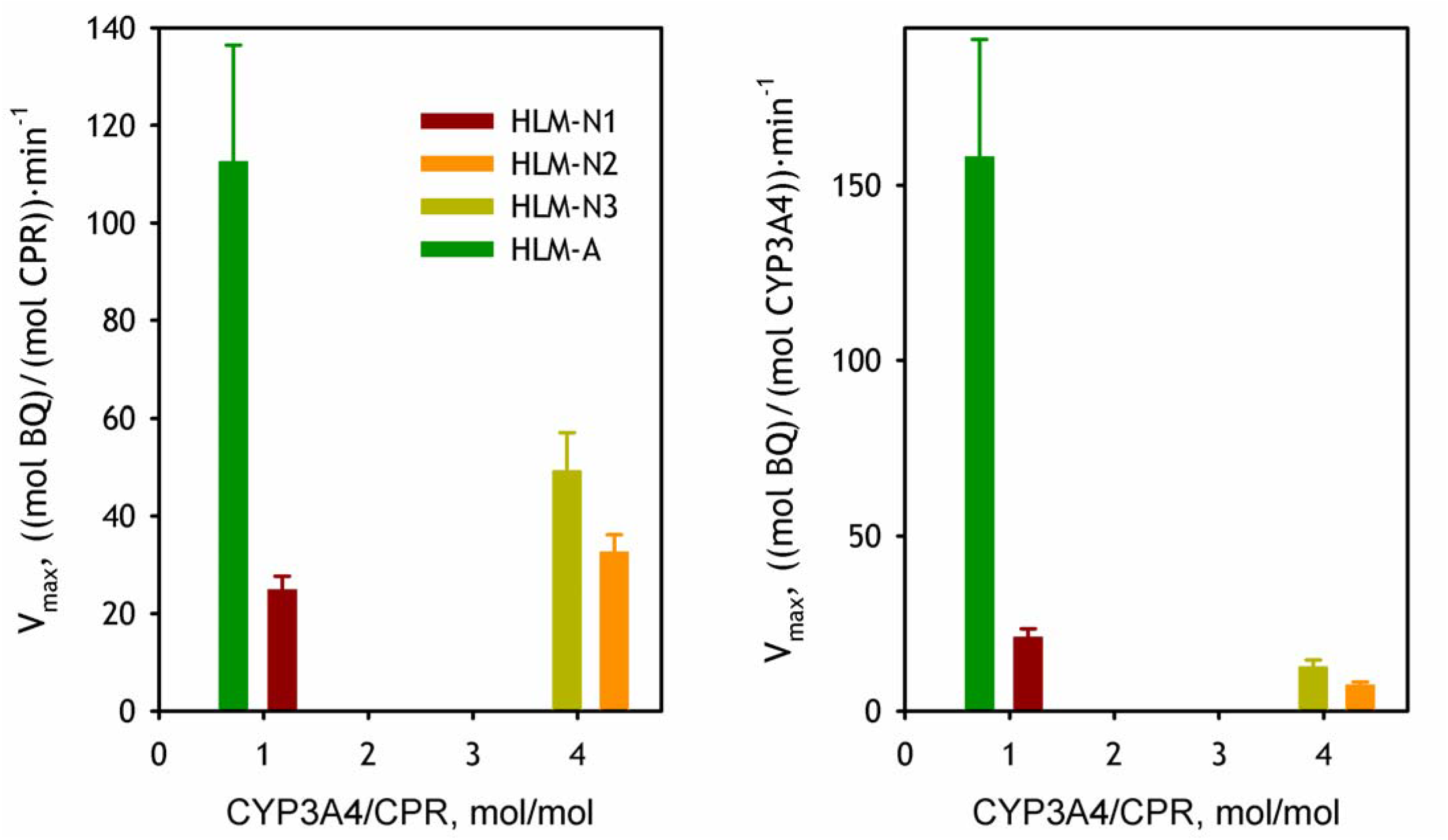
The relationship between the CYP3A4 content in HLM preparations and the rate of BQ metabolism in the presence of 25 μM ANF. The maximal velocity of BQ debenzylation is plotted against the molar ratio of CYP3A4 to CPR in microsomal membranes calculated from the results of LC-MS/MS analysis (Table 1). Left panel shows the *V*_max_ values normalized on the concentration of CPR, while the right panel shows the same results normalized per CYP3A4 content.

### The effect of ANF

Consistent with the previous observations (Domanski et al., 2001; Davydov et al., 2008; Davydov et al., 2013), the addition of 25 mM ANF completely eliminated homotropic cooperativity observed with BQ and considerably decreased the *S*_50_ values in all four HLM samples studied (Table 3). However, the effect of ANF on the maximal rate of reaction differs dramatically between the HLM samples. Whereas in the case of HLM-N1 the addition of ANF increased the maximal rate of reaction almost two times, it resulted in over two-fold inhibition of BQ metabolism in HLM-N2, while having no statistically significant effect on *V*_max_ in HLM-N3 and HLM-A (Table 3). The effect of ANF on substrate saturation curves in four HLM samples is illustrated in Fig. 1 (curves in black closed circles).

As shown in Fig. 3, where the *V*_max_ values observed in the presence of ANF are plotted against the content of CYP3A4 in HLM preparations, addition of this allosteric effector eliminates the proportionality between the rate of BQ turnover and the content of CYP3A4 in HLM-N samples. This ANF-dependent leveling of the turnover rates is achieved through the activating effect of ANF in HLM-N1, the sample with the lowest CYP3A4 content, and a substantial inhibition of BQ metabolism in CYP3A4-rich HLM-N2. At the same time, addition of ANF does not affect the contrasting difference between HLM-A and HLM-N samples in the rate of BQ turnover (Fig. 3).

### Effect of incorporation of CYP2E1 into HLM on BQ metabolism

To probe if the loss of homotropic cooperativity with BQ and a multifold increase in the rate of its metabolism in HLM-A is caused by alcohol-induced increase in the fraction of CYP2E1 in the P450 pool we studied the effect of incorporation of exogenous CYP2E1 into HLM. The dependencies of the reaction rate on substrate concentration with and without addition of ANF obtained with all four studied HLM samples enriched with CYP2E1 are exemplified in Fig. 1 (curves shown in red). The parameters of BQ metabolism in these preparations may be found in Table 3. As seen from these results, incorporation of CYP2E1 essentially eliminates the heterotropic cooperatvity in all samples studied. Furthermore, in two of the three HLM-N preparations - HLM-N1 and HLM-N3 - this incorporation results in a significant increase in the maximal rate of BQ turnover. Therefore, in some sense, the effect of incorporation of CYP2E1 into HLM on homotropic cooperativity of CYP3A4 is analogous to the effect of the addition of ANF.

Importantly, addition of ANF to CYP2E1-enriched HLM preparations largely eliminates their differences from the untreated HLM samples in the parameters of BQ metabolism. With all three CYP2E1-enriched HLM-N samples the addition of ANF considerably inhibits BQ turnover at substrate saturation, while decreasing *S*_50_ values and eliminating heterotropic cooperativity. In the case of CYP2E1-enriched HLM-A the effects of ANF on *V*_max_ and *S*_50_ are marginal, while the effect on the Hill coefficient is retained.

The effect of increasing amounts of incorporated CYP2E1 on the parameters of BQ metabolism in HLM-N and HLM-A preparations in the absence and in the presence of ANF is illustrated in Fig. 4 and Fig. 5 respectively. The dependencies of *S*_50_, *V*_max_ and the Hill coefficient on CYP2E1 content obtained in the absence of ANF reveal a striking contrast between the “normal” HLM samples and the sample obtained from the alcohol-exposed donors. Enrichment of HLM-N in CYP2E1 resulted in a stepwise decrease in the Hill coefficient, while increasing the values of *S*_50_ and *V*_max_. In contrast, incorporation of CYP2E1 into HLM-A, while also increasing *V*_max,_, had no statistically significant effect on the values of *S*_50_ and the Hill coefficient (Fig 4).

**Fig. 4.**
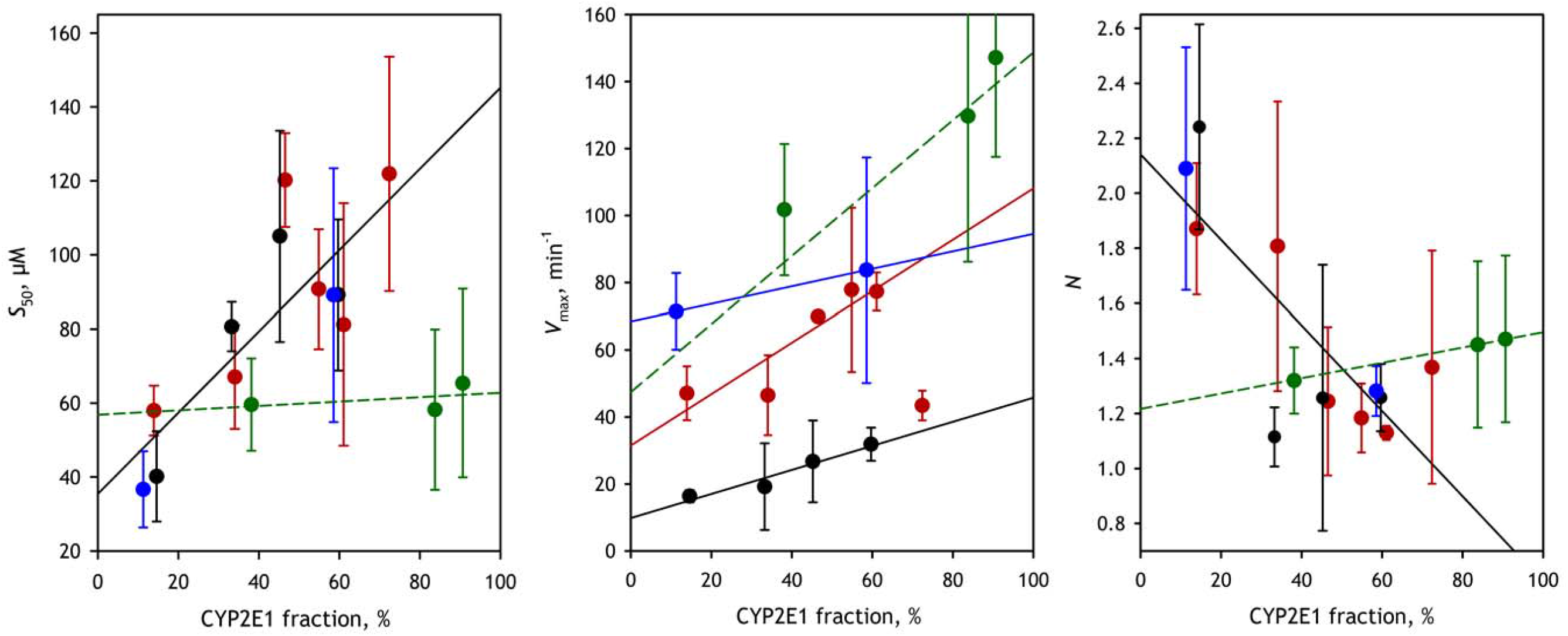
Effect of incorporation of CYP2E1 into HLM-N on the parameters of BQ metabolism at no added effector. Dependencies of *S*_50_, *V*_max_ and *N* on the fractional content of CYP2E1 in the total of 11 analyzed P450 species. The datasets obtained with HLM-N1, HLM-N2, HLM-N3 and HLM-A are shown in black, blue, red and green respectively. The error bars show the confidence intervals calculated for p=0.05. Solid lines show the linear approximations of the data.

**Fig. 5.**
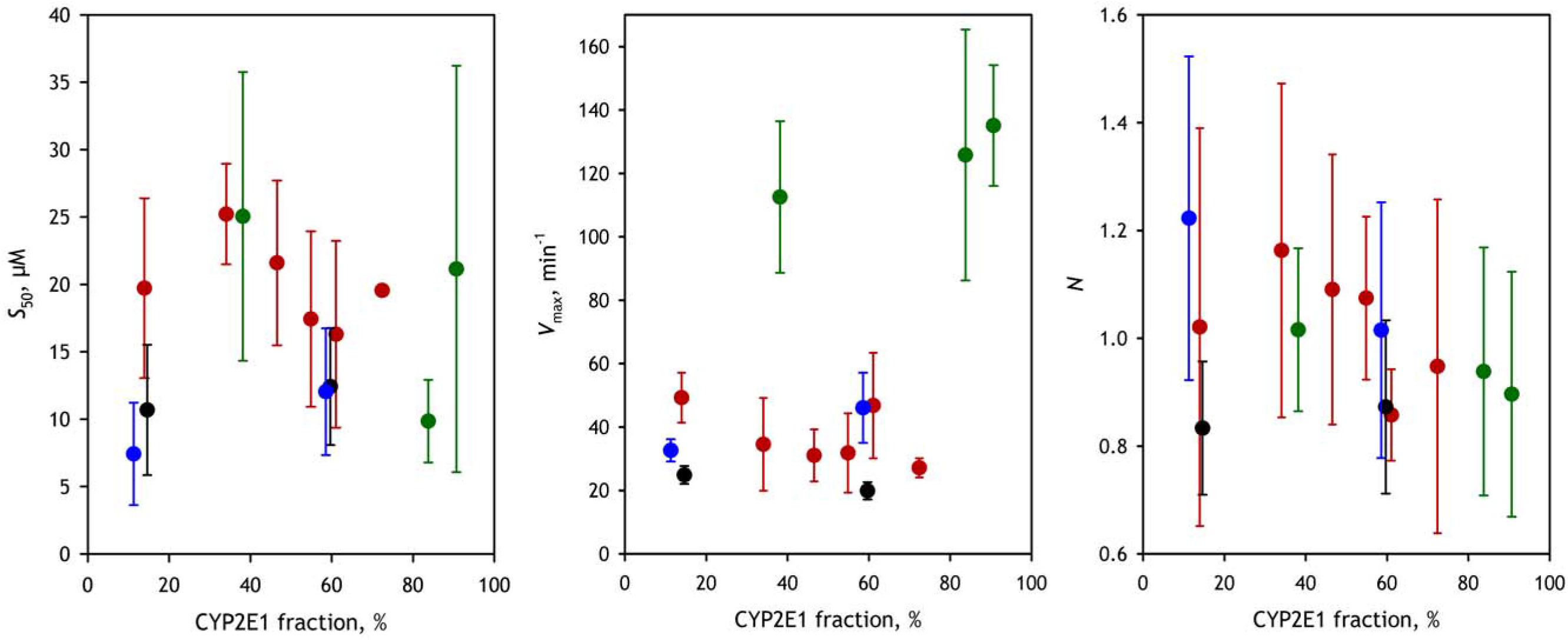
Effect of incorporation of CYP2E1 into HLM-N on the parameters of BQ metabolism in the presence of ANF. Dependencies of *S*_50_, *V*_max_ and *N* on the fractional content of CYP2E1 in the total of 11 analyzed P450 species. The datasets obtained with HLM-N1, HLM-N2, HLM-N3, and HLM-A are shown in black, blue, red and green respectively. The error bars show the confidence intervals calculated for p=0.05.

At the same time, the dependencies of the parameters of BQ metabolism on CYP2E1 content obtained in the presence of ANF (Fig. 5) show that the addition of this allosteric effector eliminates most of the effects of CYP2E1, although the increased activity of CYP2E1 in HLM-A preparation remains to be seen (Fig. 5). This observation suggests that the grounds of heterotropic cooperativity in CYP3A4 and the origins of its activation by CYP2E1 may involve some common mechanistic elements.

### Interactions of CYP2E1 with CYP3A4 in the membranes of Supersomes and their modulation by ANF

In our recent study we introduced a homo-FRET-based technique to monitor the incorporation of CYP2E1 into the microsomal membrane and probe the effect of microsomal P450 proteins on the dissociation of CYP2E1 homo-oligomers (Davydova et al., 2019). This method employs the use of CYP2E1 labeled with B0DIPY-618 maleimide in the molar ratio 1:2. Its principle is based on a deep quenching of the fluorescence of the label in the CYP2E1-B0DIPY homo-oligomers due to homo-FRET between the labels attached to different loci of the interacting protein molecules (Davydova et al., 2019). Its setup employs registration of the process of dissociation of homo-oligomers after their incorporation into the microsomal membrane through monitoring the consequent increase in the intensity of fluorescence (Fig. 5). The dependence of the amplitude of the observed increase on the amount of added microsomes (phosopholipid-to-CYP2E1-B0DIPY ratio, *R*_L/P_) allows determining the apparent dissociation constant (*K*_D_) of the homo-oligomers, or the value of *R*_L/P_ at which the degree of homo-oligomerization is equal to 50% (*R*_L/P,50%_). Effect of the presence of a given cytochrome P450 species in the microsomal membrane on *R*_L/P,50%_ is interpreted as reflecting its ability to form mixed oligomers with CYP2E1 (thus promoting the dissociatiation of the homo-oligomers) (Davydova et al., 2019).

In order to probe the relevance of the effects of CYP2E1 on the functional properties of CYP3A4 to direct physical interactions of these proteins we explored the ability of CYP2E1 to form mixed oligomers with CYP3A4 with the use of the homo-FRET based approach. In these experiments we compared SS(3A4) with the preparation of Supersomes containing recombinant CPR and cytochrome *b*_5_, but lacking any P450 protein (SS(CPR)). Dependencies of the amplitude of increase in the intensity of fluorescence on the resulting surface density of CYP2E1-B0DIPY in the membrane (or *R*_L/P_ ratio) are shown in Fig. 6. The parameters deduced by fitting these dependencies to Eq. 2 are summarized in Table 4.

**Table 4.** Effect of membrane incorporation of CYP2E1-BODIPY on the degree of its homo-oligomerization explored with homo-FRET*.

| Membranous preparation | [CYP2E1]<sub>50%</sub><sup>a</sup>,<br>pmol/cm<sup>2</sup> | (Lipid/CYP2E1)<sub>50%</sub><sup>b</sup> | FRET Efficiency, % |
| --- | --- | --- | --- |
| SS(CPR) | 0.024 ± 0.005 | 14,400 ± 2,850 | 84 ± 10 |
| SS(3A4) | 2.0 ± 0.15 | 176 ± 14 | 74 ± 9 |
| SS(3A4)+ANF | 0.58 ± 0.06 | 604 ± 62 | 84 ± 20 |

\* The values given in the Table were obtained by fitting of the titration curves with the binary association equation (Eq. 2) in combination with Eq. 3 that defines the relationship between the relative increase in the intensity of fluorescence and the steady-state concentration of the homo-oligomers. The “±” values show the confidence interval calculated for $p = 0.05$ .
<sup>a</sup> Surface density of P450 in the membranes at which the amplitude of the titration curves reaches 50% of the maximal level. This value is equal to the value of $K_D$ determined from the fitting of the titration curves with Eq. 2 combined with Eq. 3. The surface density was calculated from the $R_{L/P}$ ratio as described in Materials and Methods using Eq. 1.
<sup>b</sup> Lipid-to-P450 ratio at which the amplitude of the titration curves reaches 50% of the maximal level which was calculated from the respective [CYP2E1]<sub>50%</sub> values.
<sup>c</sup> FRET efficiency was calculated from the values of $R_{max}$ (Eq. 3) according to relationship given by Eq. 4.

**Fig. 6.**
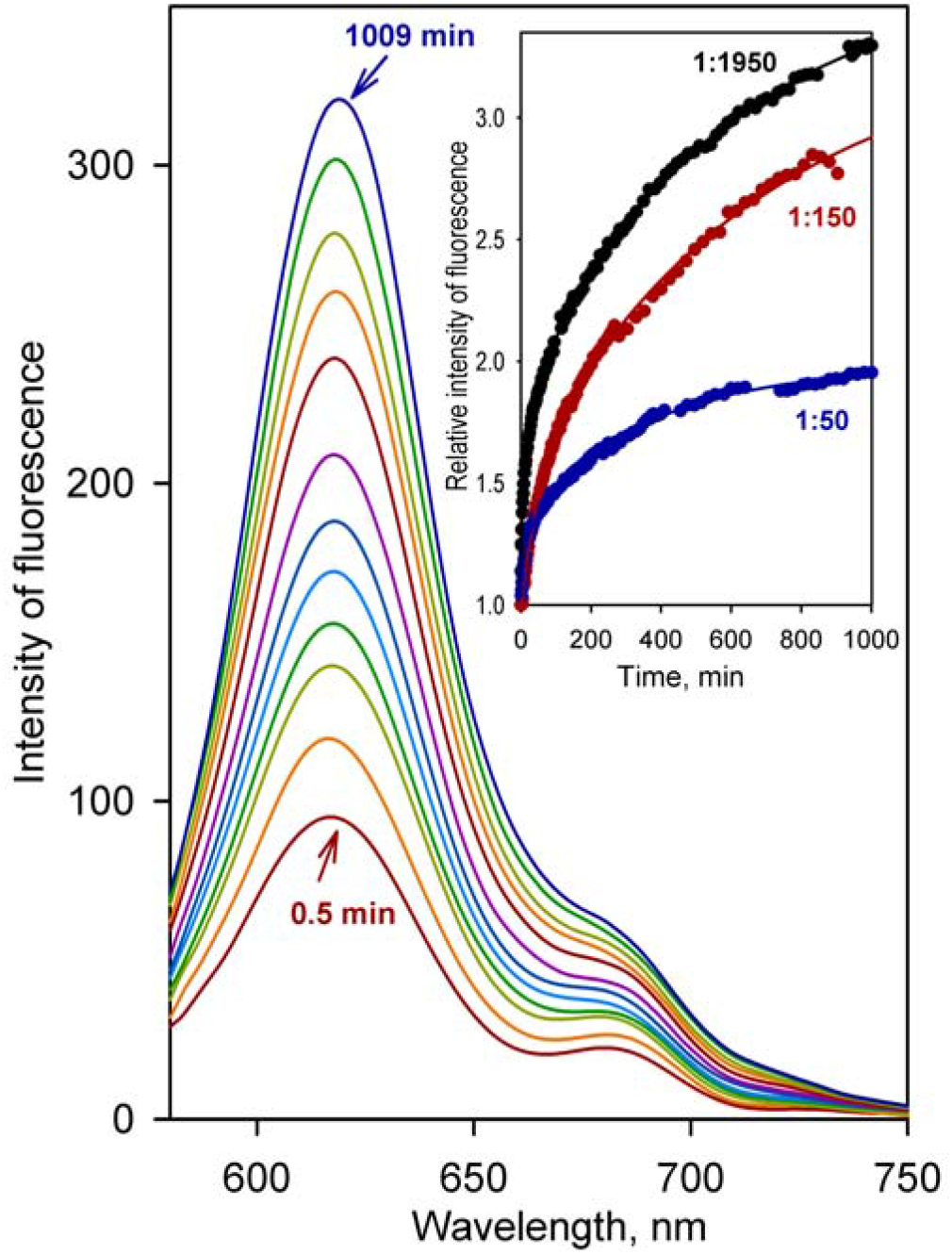
Process of dissociation of homo-oligomers of CYP2E1-BODIPY after their incorporation into microsomal membrane monitored by the changes in BODIPY fluorescence. The spectra shown in the main panel were recorded in the process of incubation of 0.081 μM CYP2E1-BODIPY with SS(3A4) added to the phospholipid concentration of 158 μM (*R*_L/P_ =1950). The spectra were recorded at 0.5, 2, 8, 16, 34, 65, 126, 253, 382, 510, 766 and 1009 min after addition of Supersomes to CYP2E1-BODIPY. Panel *c* shows the changes in the relative intensity of fluorescence (Re) during incubations performed at *R*_L/P_ ratios of 50 (blue), 150 (red) and 1950 (black). The solid lines represent the approximations of the datasets with a three-exponential equation.

Comparison of the dependencies obtained with SS(3A4) and SS(CPR) reveal a dramatic effect of microsomal CYP3A4 on homo-oligomerization of the incorporated CYP2E1-B0DIPY. If in the membranes of SS(CPR) 50% dissociation of the homo-oligomers requires the molar ratio of membrane phospholipids to CYP2E1 to be as high as 14,400:1, the presence of CYP3A4 decreases this ratio to 176:1 (Table 4). Another important finding is a two-fold decrease of the apparent maximal amplitude of the increase in fluorescence intensity in the presence of CYP3A4. This observation may reveal a conformational effect of the interactions of CYP3A4 with CYP2E1 resulting in a decreased intensity of B0DIPY fluorescence in CYP3A4-CYP2E1 complexes.

We also probed the effect of ANF on hetero-association of CYP2E1 with CYP3A4 in a series of experiments with incorporation of CYP2E1-B0DIPY into SS(3A4) in the presence of 25 μM ANF. As seen from Table 4, the addition of ANF increases the value of *R*_L/P,50%_ to 604:1 and eliminates the effect of CYP3A4 on the maximal intensity of fluorescence. These observations indicate that the addition of ANF attenuates the ability of CYP3A4 to form mixed oligomers with CYP2E1 and eliminates its apparent conformational effect revealed in decreased intensity of fluorescence of CYP2E1-B0DIPY.

## Discussion

### Changes in the composition of the cytochrome P450 ensemble under influence of chronic alcohol exposure

Analysis of the content of 11 drug-metabolizing cytochrome P450 species in pooled HLM preparation obtained from alcoholic donors revealed a 2.0 - 3.4 fold increase in the fractional content of CYP2E1 as compared to the microsomal samples unaffected by alcohol exposure (Table 1). This observation is in good agreement with the previous reports (Cederbaum, 1998; Dupont et al., 1998; Lieber, 1999; Cederbaum, 2006). 0ur data also indicate ~2 fold elevation of the fractional contents of CYP2D6 and CYP2C18 (Table 1). Although the results of our analysis of a single pooled preparation of HLM from alcohol-exposed donors cannot serve as a proof of an alcohol-induced increase in the content of these P450 species, this observation may provide a clue to further identification of changes in P450 expression induced by alcohol exposure. Importantly, our results reveal no alcohol-induced increase in the contents of CYP3A4 and CYP2A6, notwithstanding the existing reports on the induction of expression of these enzymes by ethanol in model systems (Kostrubsky et al., 1995; Feierman et al., 2003; Jin et al., 2012). This discrepancy may reveal a limited relevancy of the results obtained in model animals and cell culture to the effects of alcohol in humans. Furthermore, in the studies of the effect of alcohol on CYP3A4 expression in human hepatocytes (Kostrubsky et al., 1995) and in HepG2 cells (Feierman et al., 2003), a noticeable elevation in the CYP3A4 content was observed only upon prolonged exposure of the cells to the concentrations of ethanol of 50 mM (0.26 %) and higher, which is considerably higher than the expected physiologically-relevant concentrations of alcohol in human liver (Levitt and Levitt, 1998), at least for a long time exposure.

An intriguing and hard-to-explain peculiarity of the HLM-A preparation is a low degree of coverage of the spectrally determined quantities of both cytochrome P450 and cytochrome *b*_5_ by their amounts recovered in LC-MS/MS assays (Table 2). Since this low degree of recovery is equally characteristic to both cytochrome P450 and cytochrome *b*_5_, it is likely to stem from a low efficiency of proteolytic digestion observed with the HLM-A sample. It may reflect some characteristic features of the lipid environment in HLM-A which cause decreased accessibility of membrane-incorporated proteins for trypsinolysis.

### Multifold activation of CYP3A4 by alcohol exposure is stipulated by the effect of CYP2E1 on the landscape of P450-P450 interactions

The most striking result of this study is a demonstration of a multifold activation of CYP3A4-dependent metabolism of BQ in HLM sample from alcohol-exposed donors. Although the fractional content of CYP3A enzymes in HLM-A was the lowest among all four studied preparations, these microsomes exhibited much higher rate of metabolism and lower homotropic cooperativity with BQ as compared to HLM-N samples. Notably, in contrast to two of the three HLM-N preparations, HLM-A does not show any ANF-induced increase in *V*_max_ of debenzylation of BQ. Incorporation of additional CYP2E1 into HLM-A results in a further increase in the rate of BQ metabolism and makes this rate over an order of magnitude faster than that exhibited by the HLM-N1 sample, which is characterized by a similar CYP3A content.

The finding that the activation of BQ metabolism and attenuation of its cooperativity may be reproduced in HLM-N samples via incorporation of the exogenous CYP2E1 into their membranes suggests that these effects, at least in a part, represent direct results of the alcohol-induced increase in CYP2E1 content. It is extremely unlikely that the changes in the parameters of BQ metabolism upon CYP2E1 incorporation into HLM may result from the activity of CYP2E1 itself or be due to participation of any other P450 enzyme, as BQ metabolism is known to be almost entirely selective to CYP3A4. CYP2E1 itself lacks any potential to metabolize this substrate (Stresser et al., 2002; Davydov et al., 2015). According to the data of Stresser and co-authors (Stresser et al., 2002), the only other enzymes present in adult HLM and capable of BQ metabolism are CYP1A2 and CYP3A5. However, the turnover numbers exhibited by these enzymes with BQ are respectively 8.5 and 4.3 times lower than that characteristic to CYP3A4 (Stresser et al., 2002). Thus, due to relatively low abundance of CYP1A2 and CYP3A5 in HLM (Table 1) their sizable participation in BQ metabolism is extremely unlikely. We can conclude therefore that the effects of CYP2E1 incorporation demonstrate that the increased rate and decreased homotropic cooperativity in BQ metabolism seen in HLM-A are attributable, at least in a part, to direct effects of the increased CYP2E1 content in this sample.

The striking effects of one P450 protein on the functional properties of another P450 enzyme observed in this study and evidenced in earlier reports from our group and by others (Backes et al., 1998; Kelley et al., 2006; Reed et al., 2010; Davydov et al., 2015; Davydov et al., 2017) may be best explained with a concept of “positional heterogeneity” in P450 oligomers (Davydov, 2011; Davydov et al., 2015; Davydov, 2016; Reed and Backes, 2017; Davydova et al., 2019). This concept is based on considering heteromeric complexes of multiple P450 species as a predominant state of the P450 enzymes in the microsomal membrane. According to our hypothesis the P450 subunits forming these oligomers are not identical in their conformation and orientation, but are characterized by different abilities to be reduced, bind substrates and interact with redox partners (Davydov et al., 2015; Davydova et al., 2019). In effect, oligomerization results in abstracting an important part of the P450 pool from the catalytic activity. A discussion of possible structural basis for positional heterogeneity in P450 oligomers may be found in a comprehensive review by Reed and Backes (Reed and Backes, 2017).

From the standpoint of the above concept, the activation of one P450 enzyme by its interaction with another one might reveal a difference between the two proteins in their propensities for occupying the positions of two different types. The results of our experiments with incorporation of CYP2E1-BODIPY into CYP3A4-containing Supersomes (Fig. 7, Table 4) provide a strong evidence of high affinity interactions between CYP3A4 and CYP2E1 resulting in the formation of their heteromeric complexes. This conclusion is supported by our earlier studies with LRET-based technique (Davydov et al., 2015). It might be therefore hypothesized that, when CYP3A4 interacts with CYP2E1, the former preferentially occupies the “active” positions in the oligomers, whereas CYP2E1 fills the “restrained” ones. Resulting re-distribution of CYP3A4 between the “active” and “restrained” positions increases the active fraction of this enzyme in HLM and results in increased catalytic turnover of CYP3A4-specific substrates.

**Fig. 7.**
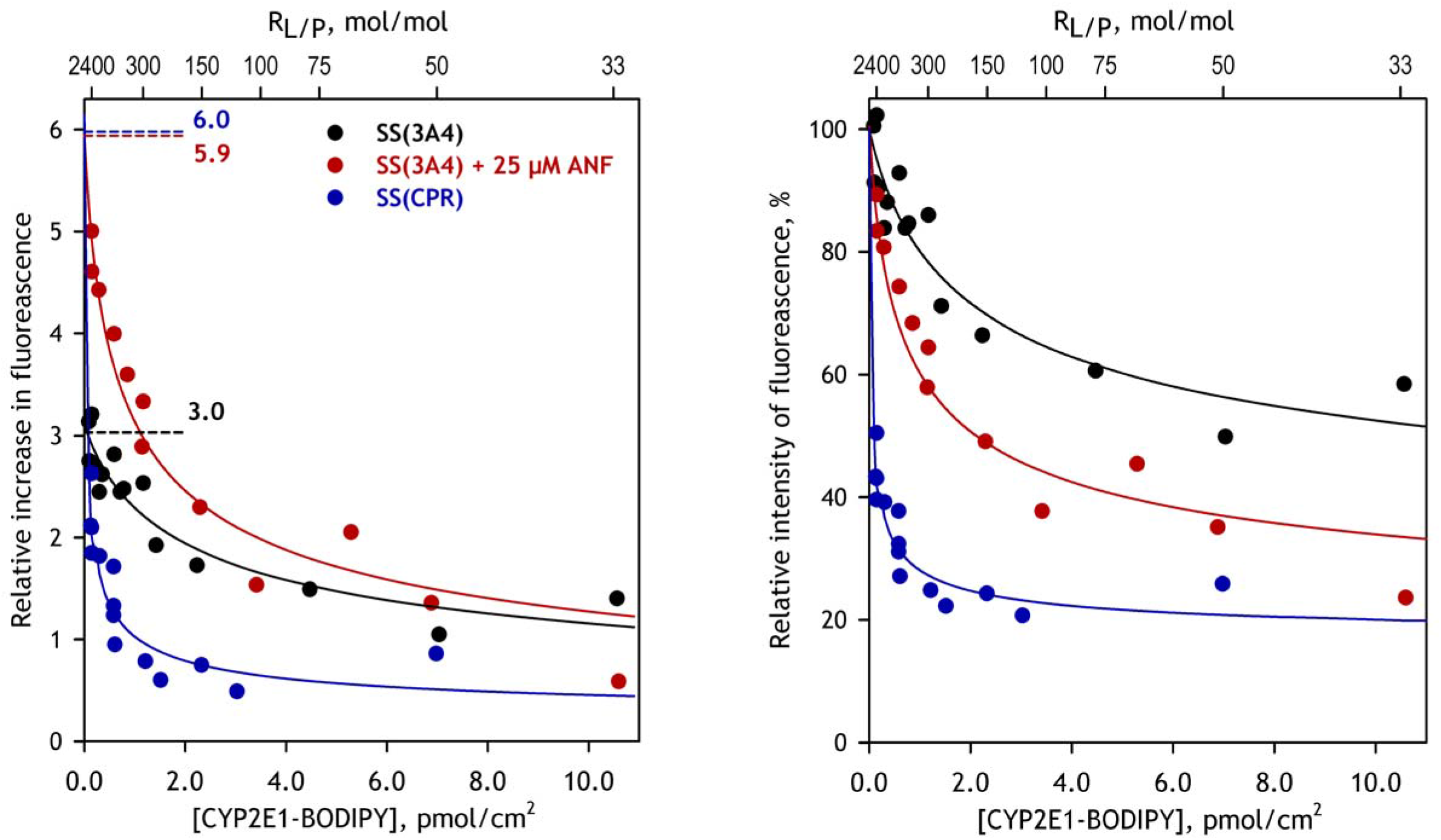
The dependencies of the changes in fluorescence caused by incorporation of CYP2E1-BODIPY into Supersomes on the concentration (surface density) of CYP2E1-BODIPY in the membrane. The datasets obtained with SS(CPR), SS(3A4) at no effector added and SS(3A4) in the presence of 25 μM ANF are shown in blue, black and red respectively. Solid lines show the approximations of the data sets with a combination of Eq. 2 and Eq. 3. Left panel shows the changes in the relative increase in the intensity of fluorescence (*R*_E_ in Eq. 3). Dashed lines shown in this panel indicate the maximal level of increase (*R*_max_) estimated from the fitting. Right panel shows the same datasets scaled to the percent of maximal fluorescence..

### Variability of heterotropic cooperativity of CYP3A4 in HLM and its modulation by P450-P450 interactions

Four preparations of HLM examined in this study revealed notable differences in their response to the addition of ANF, a prototypical allosteric effector of CYP3A enzymes. Although the attenuation of the homotropic cooperativity with BQ and increase in the affinity to this substrate were revealed with all four samples, the effect of ANF on the maximal rate of BQ turnover was contrastingly different. Whereas in the case of HLM-N1 the addition of ANF substantially increased the maximal rate of reaction, it resulted in over two-fold inhibition in HLM-N2 while having no statistically significant effect in on *V*_max_ in HLM-N2 and HLM-A (Table 3). Interestingly, these differences are correlated with the differences in the CYP3A4 content. While the activated HLM-N1 preparation has the lowest CYP3A4 content over all three HLM-N samples, the inhibited HLM-N2 preparation is characterized with the highest CYP3A4 concentration.

Importantly, increase in the fractional content of CYP2E1, either as a result of chronic alcohol exposure or through incorporation of the exogenous protein, resulted in a substantial decrease in the degree of heterotropic cooperativity seen with BQ (Table 3). Therefore, the effect of enrichment of the microsomal membrane with CYP2E1 on BQ metabolism reveals some parallelism with the effect of ANF.

The mechanisms of heterotropic cooperativity in CYP3A4 became a subject of extensive studies and a vigorous discussion over the last two decades. The initial interpretation of all known examples of CYP3A4 cooperativity, either homo- or heterotropic, is that the large substrate-binding pocket of the enzyme sometimes requires simultaneous binding of several substrate molecules to assure a productive orientation of at least one of them (see (Davydov and Halpert, 2008; Denisov et al., 2009; Denisov and Sligar, 2012) for review). The presence of at least two molecules of some substrates in the binding pocket of CYP3A4 is well established (Roberts et al., 2005; Fernando et al., 2006; Denisov et al., 2007). Although this model provides a reasonable explanation for most cases of homotropic cooperativity, it fails to explain complex instances of heterotropic activation in CYP3A4 (Davydov et al., 2013).

More recent studies compellingly demonstrated the presence of a separate allosteric effector binding site and revealed its role in the instances of heterotropic activation of CYP3A4 by various effectors, including ANF (Davydov et al., 2012; Davydov et al., 2013; Polic and Auclair, 2017; Denisov et al., 2018; Marsch et al., 2018; Denisov et al., 2019; Ducharme et al., 2019). This allosteric site is formed at the vicinity of the *F*′ and *G*′ helices and located at the interface between the two CYP3A4 molecules in the crystallographic dimer of CYP3A4 complex with peripherally-bound progesterone (PDB 1W0F (Williams et al., 2004)). As we discussed earlier (Davydov et al., 2013), the high-affinity interactions at this site require the protein to be oliogomeric - ligands are unlikely to bind with high affinity at this site in the monomeric enzyme due to small size and low degree of “buriedness” of the pocket formed by *F* - *F*′ and *G* - *G*′ loops. Therefore, if this peripheral site is involved in the mechanism of heterotropic activation of CYP3A4, the allosteric properties of the enzyme are expected to be determined by the degree of its oligomerization. A close interconnection between the allosteric properties of CYP3A4 and the degree of its oligomerization we demonstrated in our earlier study of the activating effect of ANF in model microsomes with variable surface density of CYP3A4 in the membrane (Davydov et al., 2013).

Location of the allosteric site at the interface between two subunits of the P450 oligomer suggests a critical dependence of allosteric properties of the enzyme in HLM on the formation of mixed oligomers between multiple P450 species. Although in some pairs of CYP3A4 with interacting partners this allosteric site may be retained, its affinity to the ligands and the effect of their binding on the functional properties of CYP3A4 must be considerably modified by hetero-oligomerization. Our observation that the addition of ANF attenuates the ability of CYP2E1-BODIPY to form hetero-oligomers with CYP3A4 may be interpreted as an indication of decreased ability of CYP2E1-CYP3A4 complexes to bind this allosteric effector, as compared to CYP3A4 homo-oligomers. This difference may explain the attenuation of the effect of ANF on CYP3A4 in CYP2E1-rich microsomes demonstrated in this study.

Certainly, unveiling the sources of the effect of composition of P450 ensemble on CYP3A4 allostery requires further rigorous investigation. At the current state of knowledge we are far away from understanding the mechanisms of P450 hetero-association and their relationship to the allosteric properties of the P450 ensemble. Nevertheless, the model based on the hypothesis of positional heterogeneity and the involvement of the allosteric binding site located at the interface between two subunits of P450 oligomer provides a reliable explanation for a remarkable connection between CYP3A4 allostery and the composition of the P450 pool revealed in our studies.

### Concluding remarks

Our study with human liver microsomes obtained from alcoholic donors demonstrated that chronic alcohol exposure results in a multifold activation of CYP3A4, the enzyme responsible for metabolism of wide variety of drugs currently on the market. This activation and the associated attenuation of homo- and heterotropic cooperativity of CYP3A4 can be reproduced in model HLM preparations, where the content of CYP2E1 was elevated by incorporation of the purified enzyme. These results reveal a profound effect of alcohol-induced increase in CYP2E1 content on the functional properties of CYP3A4 and suggest an important role of CYP2E1-CYP3A4 interactions in the mechanisms of the effects of alcohol exposure on drug metabolism. Furthermore, our findings compellingly demonstrate that the profile of human drug metabolism cannot be unambiguously determined from a simple superposition of the properties of the P450 species constituting the drug-metabolizing ensemble. This profile rather reflects a complex, “non-linear” combination of the functionalities of the individual enzymes which is largely affected by their intermolecular interactions and the formation heteromeric complexes.

## Acknowledgments

This research was supported by the grant R21-AA024548 from NIH. The authors are grateful to Jeffrey P. Jones (WSU) for research support, assistance in obtaining and analyzing LC-MS/MS data and continuous interest to this study. We also gratefully acknowledge the assistance of the “Human Proteome” Core Facility of the Institute of Biomedical Chemistry (IBMC, Moscow, Russia) in generating mass-spectrometry data used in characterization of HLM preparations.

## Authorship Contributions

Davydov and Zgoda designed the study; Davydov wrote the paper; Dangi, Davydova, Vavilov and Davydov conducted the experiments; Dangi, Davydov, Vavilov and Zgoda analyzed the results.

## References

Backes WL, Batie CJ, and Cawley GF (1998) Interactions among P450 enzymes when combined in reconstituted systems: formation of a 2B4-1A2 complex with a high affinity for NADPH-cytochrome P450 reductase. Biochemistry 37:12852–12859.

Bartlett GR (1959) Phosphorus assay in column chromatography. J Biol Chem 234:466–468.

Cederbaum AI (1998) Ethanol-related cytotoxicity catalyzed by CYP2E1-dependent generation of reactive oxygen intermediates in transduced HepG2 cells. Biofactors 8:93–96.

Cederbaum AI (2006) CYP2E1 - Biochemical and toxicological aspects and role in alcohol-induced liver injury. Mount Sinai J Med 73:657–672

Davydov DR (2011) Microsomal monooxygenase as a multienzyme system: the role of P450-P450 interactions. Expert Opin Drug Metab Toxicol 7:543–558.

Davydov DR (2016) Molecular Organization of the Microsomal Oxidative System: a New Connotation for an Old Term. Biochem Mosc-Suppl Ser B-Biomed Chem 10:10–21.

Davydov DR, Davydova NY, Rodgers JT, Rushmore TH, and Jones JP (2017) Toward a systems approach to the human cytochrome P450 ensemble: interactions between CYP2D6 and CYP2E1 and their functional consequences. Biochem J 474:3523–3542.

Davydov DR, Davydova NY, Sineva EV, and Halpert JR (2015) Interactions among Cytochromes P450 in Microsomal Membranes: Oligomerization of Cytochromes P450 3A4, 3A5 and 2E1 and its Functional Consequences. J BiolChem 453:219–230

Davydov DR, Davydova NY, Sineva EV, Kufareva I, and Halpert JR (2013) Pivotal role of P450-P450 interactions in CYP3A4 allostery: the case of alpha-naphthoflavone. Biochem J 453:219–230.

Davydov DR, Davydova NY, Tsalkova TN, and Halpert JR (2008) Effect of glutathione on homo- and heterotropic cooperativity in cytochrome P450 3A4. Arch Biochem Biophys 471:134–145.

Davydov DR, Deprez E, Hui Bon Hoa G, Knyushko TV, Kuznetsova GP, Koen YM, and Archakov AI (1995) High-pressure-induced transitions in microsomal cytochrome P450 2B4 in solution - evidence for conformational inhomogeneity in the oligomers. Arch Biochem Biophys 320:330–344.

Davydov DR and Halpert JR (2008) Allosteric P450 mechanisms: multiple binding sites, multiple conformers or both? Expert Opin Drug Metab Toxicol 4:1523–1535.

Davydov DR, Rumfeldt JAO, Sineva EV, Fernando H, Davydova NY, and Halpert JR (2012) Peripheral Ligand-binding Site in Cytochrome P450 3A4 Located with Fluorescence Resonance Energy Transfer (FRET). J Biol Chem 287:6797–6809.

Davydov DR, Yang ZY, Davydova N, Halpert JR, and Hubbell WL (2016) Conformational Mobility in Cytochrome P450 3A4 Explored by Pressure-Perturbation EPR Spectroscopy. Biophys J 110:1485–1498.

Davydova NY, Dangi B, Maldonado MA, Vavilov NE, Zgoda VG, and Davydov DR (2019) Toward a systems approach to cytochrome P450 ensemble: interactions of CYP2E1 with other P450 species and their impact on CYP1A2. Biochem J 476:3661–3685.

Denisov IG, Baas BJ, Grinkova YV, and Sligar SG (2007) Cooperativity in cytochrome P450 3A4 - Linkages in substrate binding, spin state, uncoupling, and product formation. J Biol Chem 282:7066–7076.

Denisov IG, Baylon JL, Grinkova YV, Tajkhorshid E, and Sligar SG (2018) Drug-Drug Interactions between Atorvastatin and Dronedarone Mediated by Monomeric CYP3A4. Biochemistry 57:805–816.

Denisov IG, Frank DJ, and Sligar SG (2009) Cooperative properties of cytochromes P450. Pharmacol Ther 124:151–167.

Denisov IG, Grinkova YV, Nandigrami P, Shekhar M, Tajkhorshid E, and Sligar SG (2019) Allosteric Interactions in Human Cytochrome P450 CYP3A4: The Role of Phenylalanine 213. Biochemistry 58:1411–1422.

Denisov IG and Sligar SG (2012) A novel type of allosteric regulation: Functional cooperativity in monomeric proteins. Arch Biochem Biophys 519:91–102.

Domanski TL, He YA, Khan KK, Roussel F, Wang QM, and Halpert JR (2001) Phenylalanine and tryptophan scanning mutagenesis of CYP3A4 substrate recognition site residues and effect on substrate oxidation and cooperativity. Biochemistry 40:10150–10160.

Ducharme J, Polic V, and Auclair K (2019) A Covalently Attached Progesterone Molecule Outcompetes the Binding of Free Progesterone at an Allosteric Site of Cytochrome P450 3A4. Bioconjugate Chem 30:1629–1635.

Dupont I, Lucas D, Clot P, Menez C, and Albano E (1998) Cytochrome P4502E1 inducibility and hydroxyethyl radical formation among alcoholics. J Hepatol 28:564–571.

Feierman DE, Melinkov Z, and Nanji AA (2003) Induction of CYP3A by ethanol in multiple in vitro and in vivo models. Alcoholism-Clinical and Experimental Research 27:981–988.

Fernando H, Halpert JR, and Davydov DR (2006) Resolution of multiple substrate binding sites in cytochrome P450 3A4: The stoichiometry of the enzyme-substrate complexes probed by FRET and Job’s titration. Biochemistry 45:4199–4209.

Gao N, Tian X, Fang Y, Zhou J, Zhang HF, Wen Q, Jia LJ, Gao J, Sun B, Wei JY, Zhang YF, Cui MZ, and Qiao HL (2016) Gene polymorphisms and contents of cytochrome P450s have only limited effects on metabolic activities in human liver microsomes. Eur J Pharm Sci 92:86–97.

Guengerich FP (1999) Cytochrome P-450 3A4: regulation and role in drug metabolism. Annu Rev Pharmacol Toxicol 39:1–17.

Jang GR and Harris RZ (2007) Drug interactions involving ethanol and alcoholic beverages. Expert Opin Drug Metab Toxicol 3:719–731.

Jin M, Kumar A, and Kumar S (2012) Ethanol-Mediated Regulation of Cytochrome P450 2A6 Expression in Monocytes: Role of Oxidative Stress-Mediated PKC/MEK/Nrf2 Pathway. Plos One 7.

Kelley RW, Cheng DM, and Backes WL (2006) Heteromeric complex formation between CYP2E1 and CYP1A2: Evidence for the involvement of electrostatic interactions. Biochemistry 45:15807–15816.

Kostrubsky VE, Strom SC, Wood SG, Wrighton SA, Sinclair PR, and Sinclair JF (1995) Ethanol and isopentanol increase CYP3A and cyp2E in primary cultures of human hepatocytes. Arch Biochem Biophys 322:516–520.

Levitt MD and Levitt DG (1998) Use of a two-compartment model to assess the pharmacokinetics of human ethanol metabolism. Alcoholism-Clinical and Experimental Research 22:1680–1688.

Lieber CS (1999) Microsomal ethanol-oxidizing system (MEOS): The first 30 years (1968-1998) - A review. Alcoholism-Clinical and Experimental Research 23:991–1007.

Marsch GA, Carlson BT, and Guengerich FP (2018) 7,8-benzoflavone binding to human cytochrome P450 3A4 reveals complex fluorescence quenching, suggesting binding at multiple protein sites. J Biomol Struct Dyn 36:841–860.

Polic V and Auclair K (2017) Allosteric Activation of Cytochrome P450 3A4 via Progesterone Bioconjugation. Bioconjugate Chem 28:885–889.

Reed J and Backes W (2016) The functional effects of physical interactions involving cytochromes P450: putative mechanisms of action and the extent of these effects in biological membranes. Drug Metab Rev 48:453–469.

Reed JR and Backes WL (2012) Formation of P450·P450 complexes and their effect on P450 function. Pharm Ther 133:299–310.

Reed JR and Backes WL (2017) Physical Studies of P450-P450 Interactions: Predicting Quaternary Structures of P450 Complexes in Membranes from Their X-ray Crystal Structures. Front Pharmacol 8.

Reed JR, Eyer M, and Backes WL (2010) Functional interactions between cytochromes P450 1A2 and 2B4 require both enzymes to reside in the same phospholipid vesicle. Evidence for physical complex formation. J Biol Chem 285:8942–8952.

Roberts AG, Campbell AP, and Atkins WM (2005) The thermodynamic landscape of testosterone binding to cytochrome P450 3A4: ligand binding and spin state equilibria. Biochemistry 44:1353–1366.

Smith PK, Krohn RI, Hermanson GT, Mallia AK, Gartner FH, Provenzano MD, Fujimoto EK, Goeke NM, Olson BJ, and Klenk DC (1985) Measurement of protein using bicinchoninic acid. Anal Biochem 150:76–85.

Spatzenegger M, Liu H, Wang QM, Debarber A, Koop DR, and Halpert JR (2003) Analysis of differential substrate selectivities of CYP2B6 and CYP2E1 by site-directed mutagenesis and molecular modeling. J Pharm Exp Ther 304:477–487.

Stresser DM, Turner SD, Blanchard AP, Miller VP, and Crespi CL (2002) Cytochrome P450 fluorometric substrates: Identification of isoform-selective probes for rat CYP2D2 and human CYP3A4. Drug Metabolism and Disposition 30:845–852.

Wang X, He B, Shi J, Li Q, and Zhu H-J (2020) Comparative Proteomics Analysis of Human Liver Microsomes and S9 Fractions. Drug Metabolism and Disposition 48:31.

Wibo M, Amar-Costesec A, Berthet J, and Beayfay H (1971) Electron microscope examination of subcellular fractions III. Quantitative analysis of the microsomal fraction isolated from rat liver. J Cell Biol 51:52–71.

Williams PA, Cosme J, Vinkovic DM, Ward A, Angove HC, Day PJ, Vonrhein C, Tickle IJ, and Jhoti H (2004) Crystal structures of human cytochrome P450 3A4 bound to metyrapone and progesterone. Science 305:683–686.

Zhang HF, Wang HH, Gao N, Wei JY, Tian X, Zhao Y, Fang Y, Zhou J, Wen Q, Gao J, Zhang YJ, Qian XH, and Qiao HL (2016) Physiological Content and Intrinsic Activities of 10 Cytochrome P450 Isoforms in Human Normal Liver Microsomes. J Pharm Exp Ther 358:83–93.

